# callsync: an R package for alignment and analysis of multi-microphone animal recordings

**DOI:** 10.1101/2023.02.07.527470

**Authors:** Simeon Q. Smeele, Stephen A. Tyndel, Barbara C. Klump, Gustavo Alarcón-Nieto, Lucy M. Aplin

## Abstract

1. To better understand how vocalisations are used during interactions of multiple individuals, studies are increasingly deploying on-board devices with a microphone on each animal. The resulting recordings are extremely challenging to analyse, since microphone clocks drift non-linearly and record the vocalisations of non-focal individuals as well as noise.
2. Here we address this issue with `callsync`, an R package designed to align recordings, detect and assign vocalisations to the caller, trace the fundamental frequency, filter out noise and perform basic analysis on the resulting clips.
3. We present a case study where the pipeline is used on a dataset of six captive cockatiels (*Nymphicus hollandicus*) wearing backpack microphones. Recordings initially had drift of ∼2 minutes, but were aligned to within ∼2 seconds with our package. Using `callsync`, we detected and assigned 2101 calls across three multi-hour recording sessions. Two had loud beep markers in the background designed to help the manual alignment process. One contained no obvious markers, in order to demonstrate that markers were not necessary to obtain optimal alignment. We then used a function that traces the fundamental frequency and applied spectrographic cross correlation to show a possible analytical pipeline where vocal similarity is visually assessed.
4. The `callsync` package can be used to go from raw recordings to a clean dataset of features. The package is designed to be modular and allows users to replace functions as they wish. We also discuss the challenges that might be faced in each step and how the available literature can provide alternatives for each step.

## Introduction

The study of vocal signals in animals is a critical tool for understanding the evolution of vocal communication (Endler 1993). Vocalisations commonly occur in group contexts, where they may function to signal movement or identity, and mediate interactions. However, the problem of assigning identity has led most previous work to focus on long-range calls or on vocalisations made when alone, e.g., territorial calls and song, and to focus on recording one individual at a time. Yet only by studying the ways that animals communicate in ‘real time’ can allow us to untangle the complicated dynamics of how group members signal one another (Gill et al. 2015). For example, the context of how individuals address members of their social network (Cheney and Seyfarth 2018), and the ensuing call and response-dynamics (Araya-Salas et al. 2020), can only be studied by recording multiple individuals simultaneously. These communication networks can help us understand how animals coordinate call-response with movement (Demartsev et al. 2023) as well as the function of vocal imitation (Dahlin et al. 2014; Knörnschild et al. 2012; Nousek et al. 2006).

Recent innovations in recording technologies have allowed for a dramatic increase of fine scale on-animal bioacoustic data collection (Wild et al. 2022; Bravo Sanchez et al. 2021; Gill et al. 2016), allowing multiple individuals to be recorded simultaneously. However, as the capability of placing small recording devices on animals increases, so does the need for tools to process the resulting data streams. Several publicly available R packages exist that measure acoustic parameters from single audio tracks (*seewave*: Sueur, Aubin, and Simonis (2008), *tuneR*: Ligges et al. (2022), *WarbleR*: Araya-Salas and Smith-Vidaurre (2017)), but to our knowledge, none addresses the critical issue of microphone clock drift and the ability to align and process multiple recordings. This poses a serious issue for those studying communication networks of multiple tagged individuals. In this paper, we apply a new R package, `callsync`, that aligns multiple misaligned audio files, detects vocalisations, assigns these to the vocalising individual and provides an analytical pipeline for the resulting synchronised data.

The primary target for use of this package are researchers that study animal communication systems within groups. As researchers deploy multiple microphones that record simultaneously, the resulting clock drift can prove to be a barrier in further data processing steps (Schmid et al. 2010). To make matters worse this drift is often non-linear (Anisimov et al. 2014). Thus, if several microphone recorders (whales; Miller and Dawson (2009), Hayes et al. (2000), bats; Stidsholt et al. (2019)) are placed on animals, it is critical for researchers to be able to line up all tracks so that calls can be assigned correctly to the vocalising individual (loudest track). The main functionality of `callsync` is to align audio tracks, detect calls from each track, determine which individual (even ones in relatively close proximity to one another) is vocalising and segment them (detect start and end of signal as in detection module), as well as take measurements of the given calls (see Fig. 1). As `callsync` takes a modular approach to aligning, detecting, and analysing audio tracks, researchers can use only the components of the package that suit their needs.

**Figure 1:**
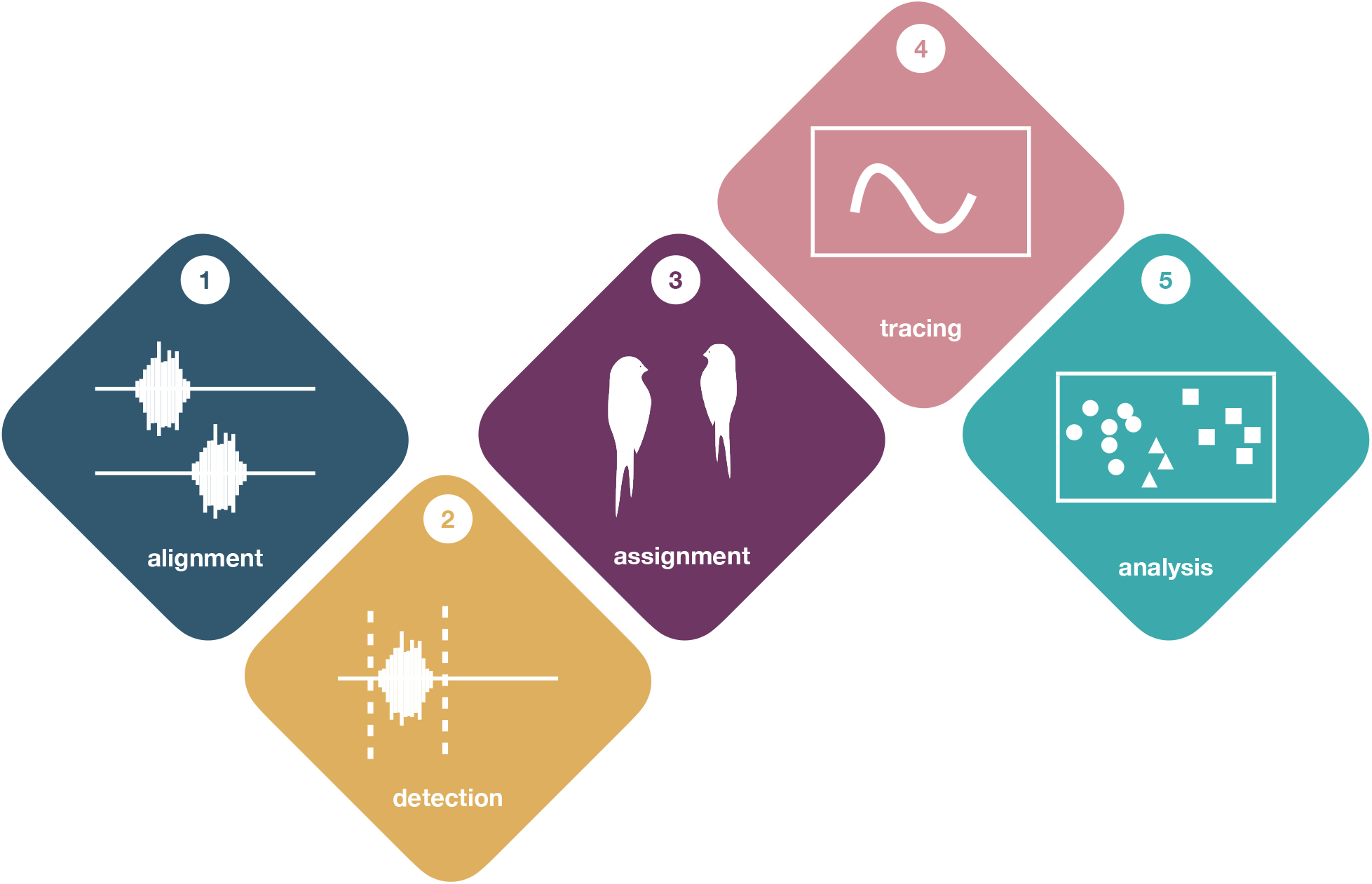
Flowchart from the `callsync` package. (1) The *alignment* module can be used to align multiple microphones that have non-linear temporal drift. (2) The *detection* module can be used to detect vocalisations in each recording. Filters can be applied to remove false alarms in the detection module. (3) The *assignment* module can be used to assign a vocalisation to the focal individual, making sure that vocalisations from conspecifics are excluded from the focal recording. (4) The *tracing* module can be used to trace and analyse the fundamental frequency for each vocalisation. (5) The final *analysis* module can be used to run spectrographic cross correlation and create a feature vector to compare across recordings.

Current research packages that implement call alignment strategies are either used in Matlab (Malinka et al. 2020; Anisimov et al. 2014) or c++ (Gill et al. 2015). However, these tools have not, up to now, been adapted for the R environment, a popular programming language among many animal behaviour and bioacoustic researchers. Many of these tools are not documented publicly nor open source, and can require high licensing fees (i.e., Matlab). While the design of our package is best suited to contexts where all microphones exist in the same spatial area, it is the goal that it can be adapted to other contexts. `callsync` is publicly available on CRAN and GitHub, is beginner friendly with strong documentation, and does not require extensive programming background. This open-source tool will allow researchers to expand the study of bioacoustics and solve an issue that impedes detailed analysis of group-level calls.

### Generate example: Spring Peeper

We downloaded two Spring Peeper (Pseudacris crucifer) recordings from the Macaulay Library at the Cornell Lab of Ornithology (ML397067 and ML273028). Using these two recordings, we generated two artificial audio tracks, one drifted approximately 18 seconds from the other. Drift was simulated by inserting 0.03s of background noise every second in one of the tracks. Background noise for each track was the same, and constructed by concatenating random snippets of noise from each of the Macaulay Library recordings. The background noise was then amplified by 4dB.

Both artificial tracks incorporated 10 calls from each Macaulay recording. The calls were arranged to simulate alternating calling behaviour between the two frogs. Using the *tuneR* package (Ligges et al. 2022), we normalised the calls to simulate the presence of a focal individual, setting the focal calls to 85% and background calls to 55% of their maximum amplitude, inversely for each track.

We then used `callsync` to align the tracks using 2 minute chunks, and showed that all detected calls were assigned to the correct individual. We also showed that the fundamental frequency could be correctly traced. For details see the vignette in the CRAN version of `callsync`.

### Case study: cockatiels

We present a case study to show how `callsync` functions can be included in a workflow (see Fig. 1). We used a dataset of domestic cockatiels (*Nymphicus hollandicus*). These birds are part of an ongoing study at the Max Planck Institute of Animal Behavior in Radolfzell, Germany. 30 birds were housed in five groups of six individuals, with each group of six housed separately in a 4x3x2.7m aviary facility. Each bird was fitted with a TS-systems EDIC-Mini E77 tag inside a sewn nylon backpack fitted via Teflon harness around the wings, with the total weight of all components under 7% of body weight (weight range of birds 85-120g). Audio recordings were scheduled to record for 4 hours per day. Each microphone was automatically programmed to turn on and off daily at the same time. For the purposes of demonstration, three full recording sessions (ca. four hours) were selected for processing where the microphones were scheduled to record starting at approximately sunrise (July 15th-16th, 2021 and December 08, 2022). Two of the recording sessions (2021) included manually played beeps in the background; one every hour at 10khz, one every 10 minutes at 0.4khz. This was done to ensure that the alignment would work both with and without external assistance, as some circumstances for research might allow such an inclusion, while others may prohibit it. Seven days after deployment, microphone recorders were removed and recordings were downloaded as .wav files directly onto a computer from the tag, according to the manufacturer protocols. Data from each microphone were placed into the appropriate folder (see workflow instructions) and processed in accordance with our package workflow.

### Installation and set-up

The package call be installed from CRAN running `install.packages(‘callsync’)` or a developmental version can be installed from GitHub (https://github.com/simeonqs/callsync):

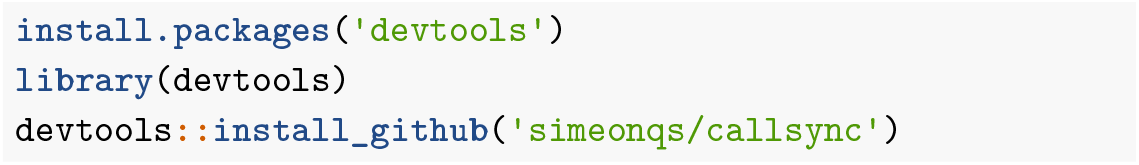

While all underlying code and functions from the CRAN repository can be found in the aforementioned GitHub repository, we have created a separate repository from which readers can follow the code from the case study: https://github.com/simeonqs/callsync_an_R_package_for_alignment_and_analysis_of_multi-microphone_animal_recordings. All required packages are also automatically installed and loaded for the case study when running the `00_set_up.R` script from this repository.

### Alignment of raw recordings

The general goal of the alignment step is to shift unaligned recordings to a degree that will allow vocalisations to be processed further. While researchers’ requirements on alignment precision will vary, the output of this function should allow researchers to be able to identify that vocalisations picked up across all microphones are identified as the same call off of every microphone. It should be noted that recordings are still not perfectly aligned, since clock-drift occurs within the chunks.

The raw recordings consisted of three wav files of ca. four hours for six cockatiels (i.e., 18 files total). The backpack microphones have internal clocks that sync with the computer clock when connected via USB and automatically turn recordings on and off. However, these clocks drift in time both during the off period, creating start times that differ up to a few minutes, and also during the recording period, causing additional variable drift up to a minute in four hours of recording for the microphone model we used. The function `align` can be used as a first step to align the audio recordings. To accurately align these tracks, the full audio files were automatically split into 15 minute chunks of recording to ensure that drift was reduced to mere seconds. This value can be adjusted depending on the amount of drift. The function selects one recording (focal recording) and aligns all the other recordings relative to the selected one recording using cross correlation on the energy content (summed absolute amplitude) per time bin (function step_size), in our case 0.5 seconds was used. This value can also be adjusted, lower values allow for greater precision but higher computational load. The call detection and assignment step (see below) includes fine alignment and we therefore accepted a potential error of 0.5 seconds. There is also an option to store a log file that records the offset for each chunk.

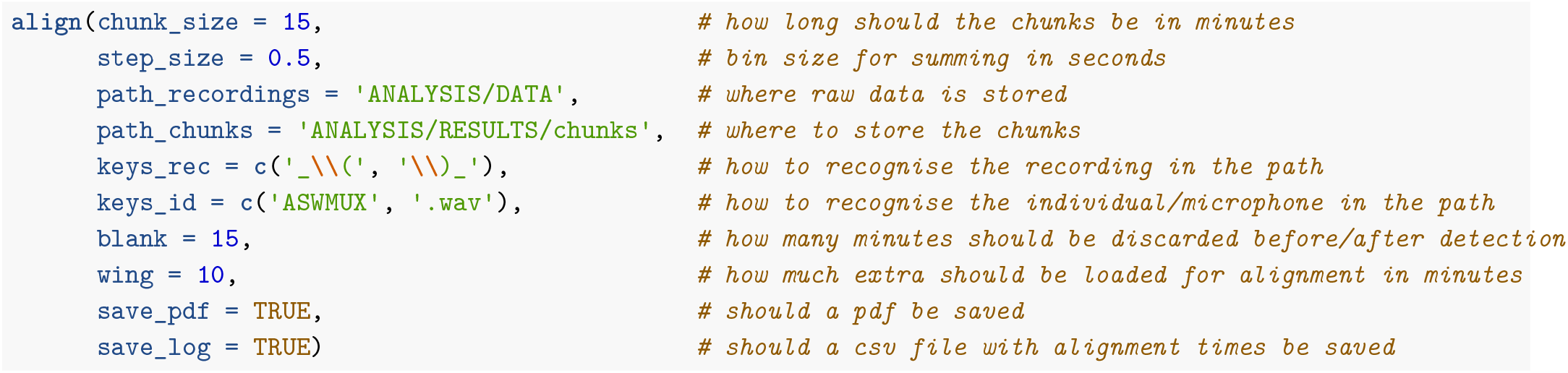

For cross correlation, the function `align` loads the chunks with additional minutes before and after (option `wing`) to ensure that overlap can be found. The cross correlation is performed using the function `simple.cc`, which takes two vectors (the binned absolute amplitude of the two recordings) and calculates the absolute difference while sliding the two vectors over each other. It returns the position of minimum summed difference, or in other words, the position of maximal overlap. This position is then used to align the recordings relative to the first recording and save chunks that are maximally aligned. Note that due to drift during the recording, the start and end times might still be seconds off; it is the overall alignment of the chunks that is optimised. The function also allows the user to create a pdf document with waveforms of each individual recording and a single page per chunk (Fig. 2), to visually verify if alignment was successful. For our dataset all chunks but one aligned correctly without a filter. If this is not the case the option `ffilter_from` can be set to apply a high-pass filter to improve alignment. Mis-aligned chunks can also be rerun individually using the chunk size argument (argument chunk_seq) to avoid re-running the entire dataset. This was done for the case study as well (recording session from 16th July 2021, start time 15 minutes).

**Figure 2:**
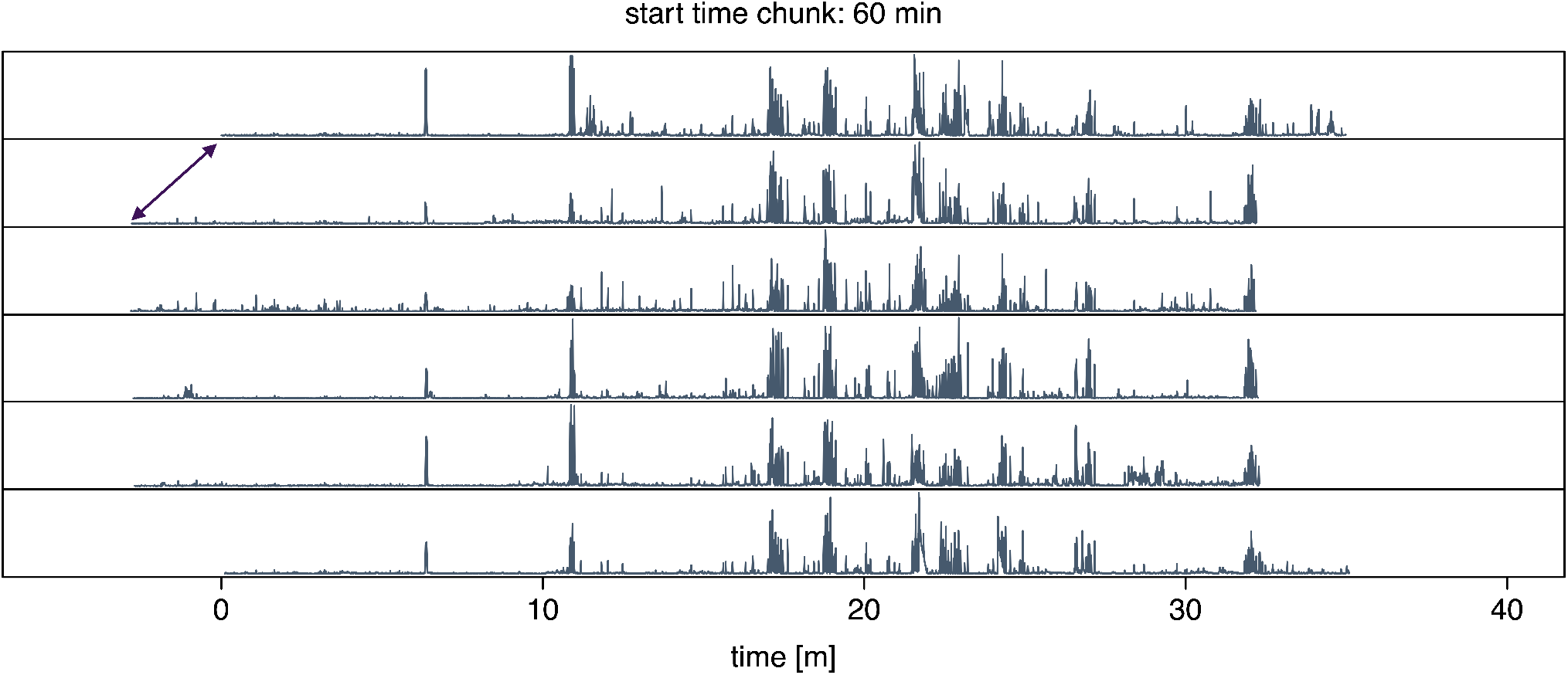
Example of the alignment output. Olive-coloured lines represent the summed absolute amplitude per bin (= 0.5 seconds). Recordings are aligned relative to the first recording (which starts at 0). Note that recordings 2-5 initially started ∼2 minutes earlier (purple arrow) but are now aligned. The title displays the start time of the chunk in the raw recording.

### Call detection and assignment

The next step is to detect calls in each set of chunks and assign them to the correct source individual. The `detect.and.assign` function wrapper loads the chunks using the function `load.wave` where it optionally applies a high-pass filter to reduce the amount of low frequency noise. To detect calls, ‘detect.and.assign’ first calls the function `call.detect.multiple`, which is used to detect multiple calls in an R wave object. It first applies the `env` function from the *seewave* (Sueur, Aubin, and Simonis 2008) package to create a smooth Hilbert amplitude envelope. It then detects all the points on the envelope which are above a certain threshold relative to the maximum of the envelope. After removing detections that are shorter than a set minimum duration (argument `min_dur`) or longer than a set maximum (argument `max_dur`) it returns all the start and end times as a data frame.

Because the microphones on focal individuals are very likely to record the calls of the non-focal individuals as well, we implemented a step that assigns the detected calls to the individual emitting the sound, based on amplitude. For this, `detect.and.assign` subsequently calls the function `call.assign`, which runs through all the detections in a given chunk for a given individual (i.e., output of `call.detect.multiple`) and runs the `call.detect` function to more precisely determine the start and end time of the call. It then ensures that minor temporal drift after the align function is corrected by rerunning the `simple.cc` function pairwise between the focal recording and all others. After alignment it calculates the summed absolute energy content on all recordings for the time frame when the call was detected and compares the recording where the detection was made to all other recordings. If the loudest recording is louder by a set percentage (this value can be adjustable according to the researchers needs) than the second loudest recording, the detection is saved as a separate wav file. If not, this means it’s not possible to determine the focal individual and the detection is discarded (there is an option to save all detections before the assignment step). The function also allows the user to create a pdf file with all the detections (see Fig. 3 for a short example) to manually inspect the results.

**Figure 3:**
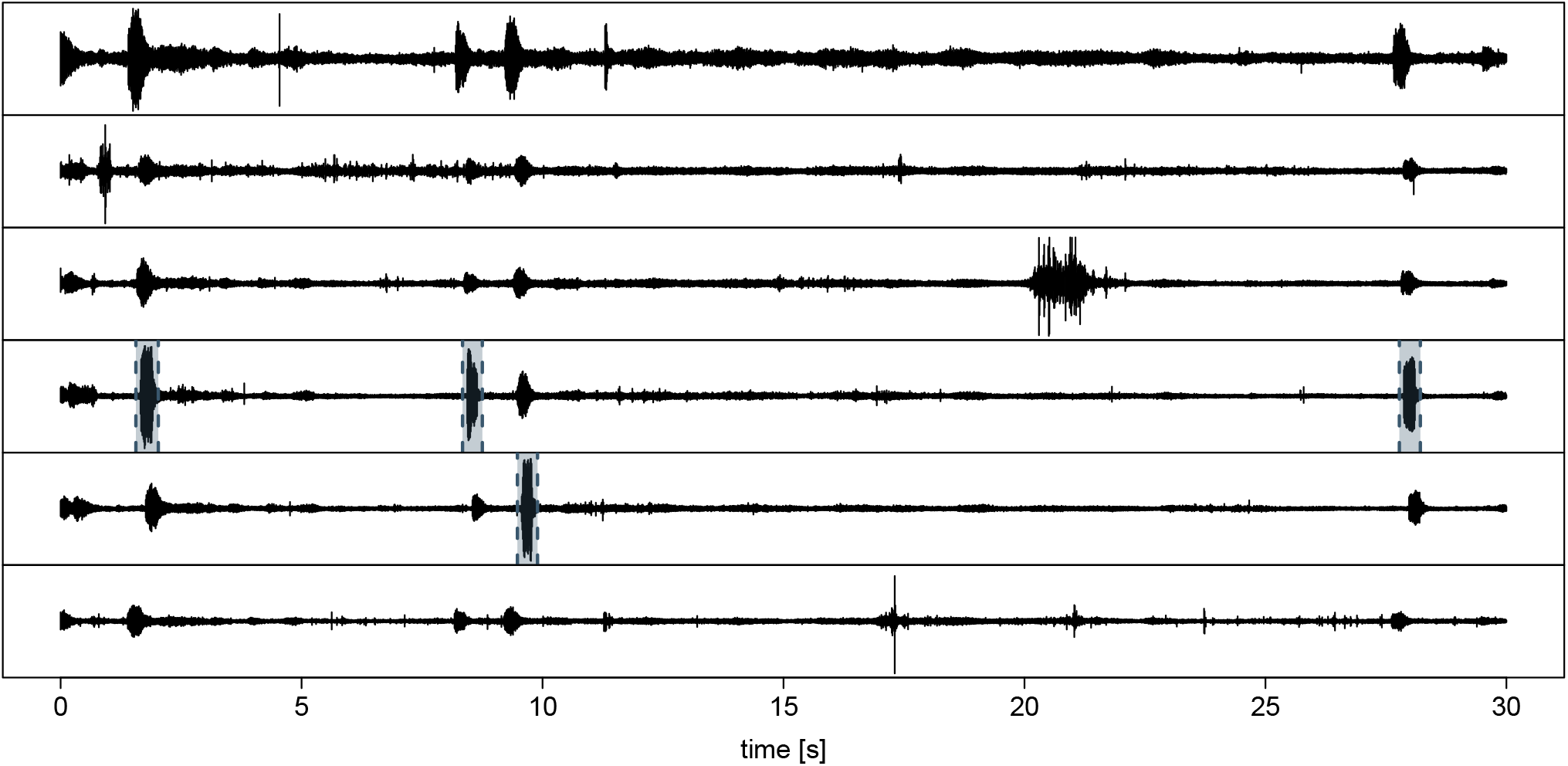
Example of the detection output. Black lines are the waveforms. Olive-coloured dashed lines with shaded areas in between are the detected calls. Note that only the loudest version of each call is selected. The first call is also loud in the first channel (top-row) but contains less energy throughout the whole call (channel four is loud for a longer duration). Animals with individual specific-sounds (scratching, etc) are seen clearly on only single microphones.

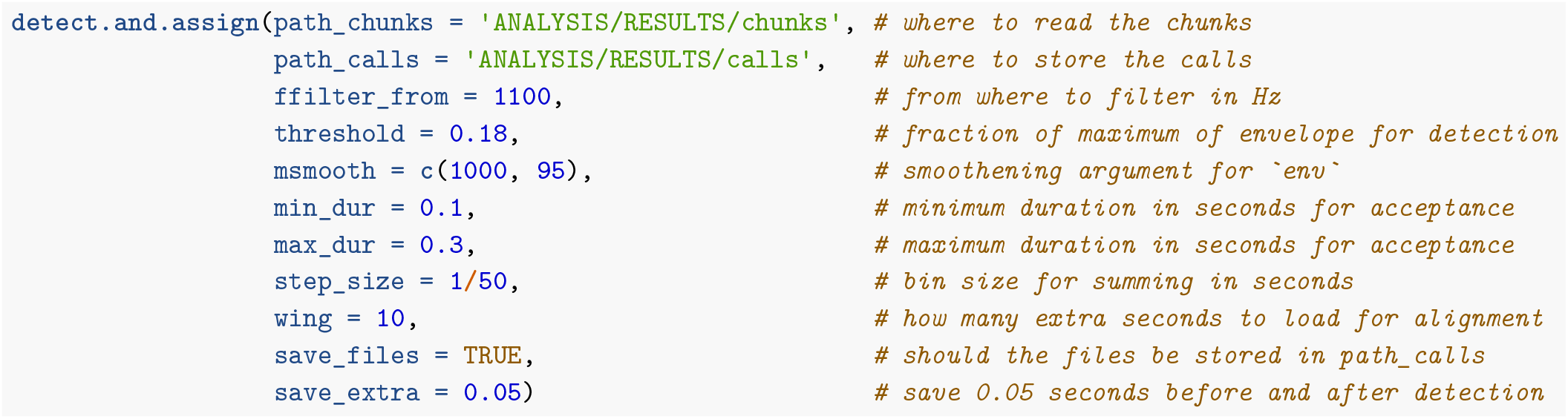

For the 12-hour long cockatiel dataset, the function detected and assigned 4409 events, 2101 of which were retained as vocalisations (see next step). The removed events were primarily noise and noisy calls (e.g., overlapping calls and calls with echo). To estimate the accuracy of the function we manually detected and assigned IDs for 300 calls in three chunks (chosen due to high vocal activity), ran the same filtering step as `callsync` functions described earlier to get rid of 100 noisy examples, and compared the performance of the `detect.and.assign` function to manually labelled data. The ground truth was performed using the function `calc.perf`. This function simply determines whether the detected calls (and subsequent labelled ids) of two datasets match. If sufficient temporal overlap exists between the two datasets, they are determined to match. The false positive rate was <4%, with noise (two detections) or noisy calls (four detections) being responsible; and the true positive rate was 81%. The remaining 19% (false negatives) were mostly calls that were too quiet to be picked up by our choosen amplitude threshold.

## Tracing

To analyse the calls, the chunks were loaded and the function `call.detect` was run to determine the start and end times of the call. The wave objects were then resized to only include the call (`new_wave`). To trace the fundamental frequency we applied the `trace.fund` function to the resized wave objects. This function detects the fundamental frequency in a sliding window based on the spectrum and a relative threshold. We ran the latter step in parallel using the function `mclapply` from the base R package *parallel* (see Fig. 4a for an example).

**Figure 4:**
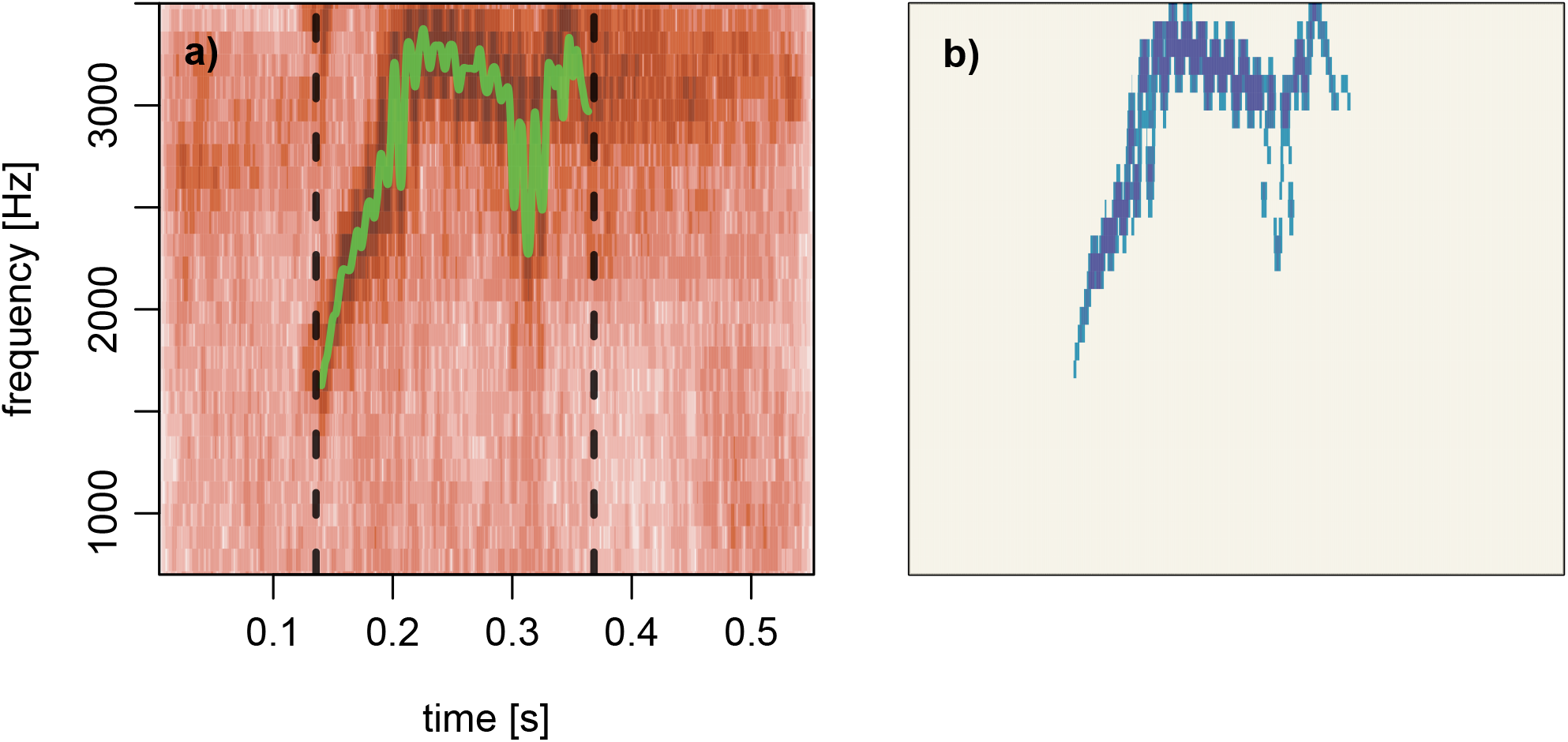
a) Spectrogram of a cockatiel call with start and end (black dashed lines) and the fundamental frequency trace (green solid line). b) Noise reduced spectrogram where darker colours indicate higher intensity.

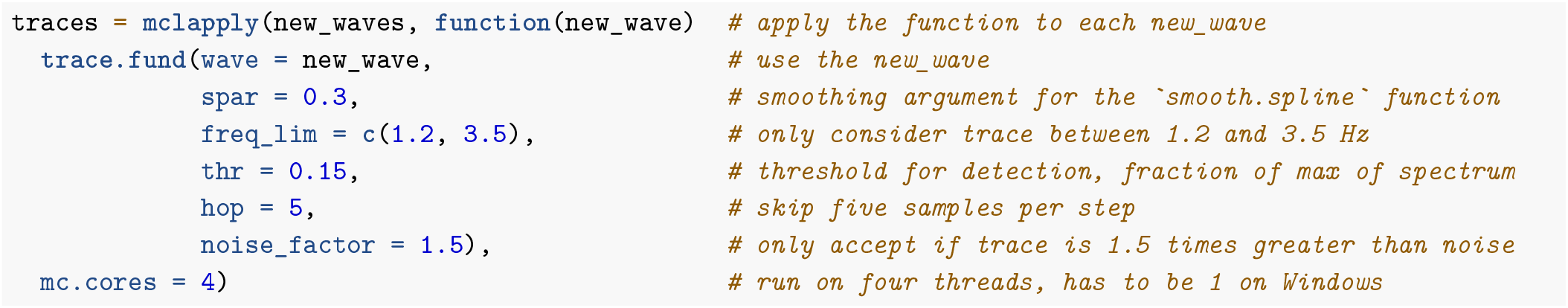

Since the call detection step also picks up on a lot of noise (birds scratching, flying, walking around) as well as calls, we ran a final step to filter the measurements and traces before these were saved.

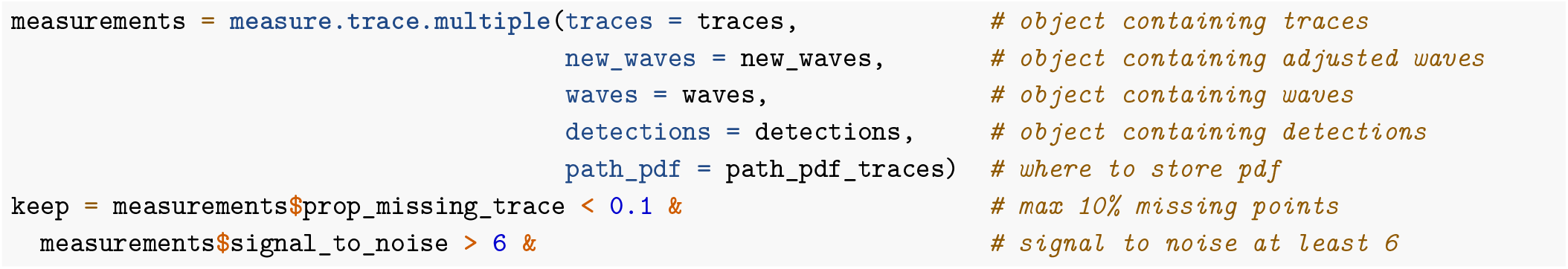

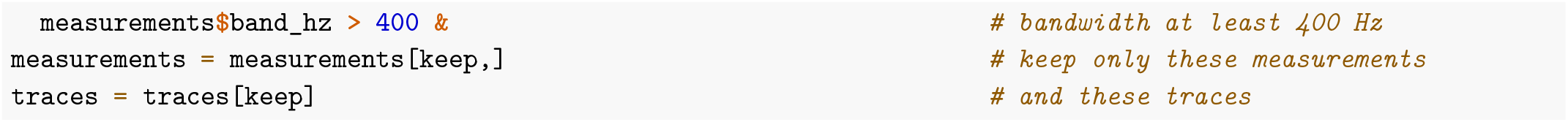

## Analysis

A frequently used method to compare calls is to measure their similarity using spectrographic cross correlation (SPCC) (Cortopassi and Bradbury 2000), where two spectrograms are slid over each other and the pixelwise difference is computed for each step. At the point where the signals maximally overlap one will find the minimal difference. This score is then used as a measure of acoustic distance between two calls. The function `run.spcc` runs SPCC and includes several methods to reduce noise in the spectrogram before running cross correlation (for an example see Fig. 4b). To visualise the resulting feature vector from running SPCC on the cockatiel calls we used uniform manifold approximation and projection (UMAP) (Konopka 2022) which projects the results in two-dimensional space. Calls cluster very strongly by the two separate recording groups (2021 and 2022), giving some evidence of different vocal signatures (see Fig. 5). This is a very simple analysis, and we only included it to illustrate a potential use of the results from the previous steps.

**Figure 5:**
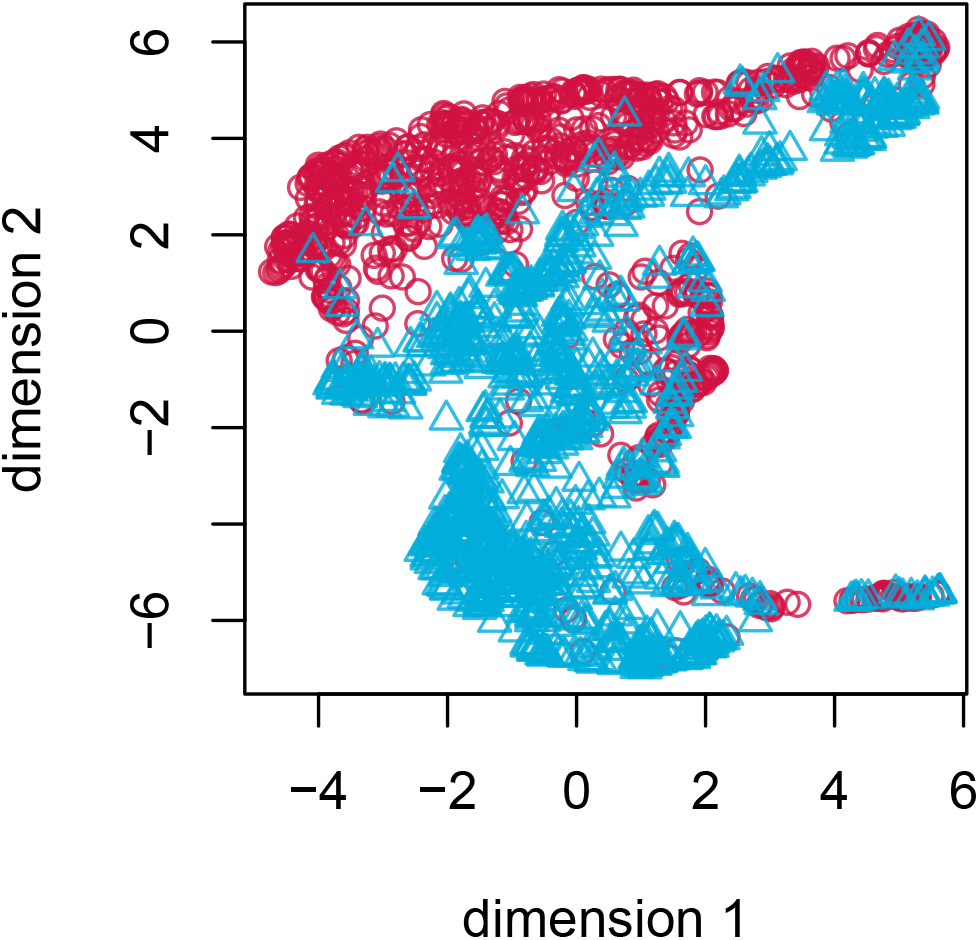
Call distribution in uniform manifold approximation and projection space. Dots represents calls and are coloured by year.

## Discussion

Here we detail our R package `callsync`, designed to take raw microphone recordings collected simultaneously from multiple individuals and align, extract and analyse their calls. We present callsync performance on a computer generated dataset of 10 minutes of recording from two frogs and a case study of 12 hours of natural recordings from two times six communally housed captive cockatiels. Each of the modular components (alignment, detection, assignment, tracing, and analysis) successfully achieved the stated goals in both systems. In the computer generated dataset, no errors were made. In the case study, misaligned audio tracks were accurately aligned in a first step (see Fig. 2), calls were correctly identified in the aligned recordings (see Fig. 3), the individual making the call was selected (see Fig. 3), and downstream data analysis was performed (Fig. 4, Fig. 5). `callsync` can perform alignment even on inter-microphone drift that constitutes minutes as well as handle unpredictable and non-linear drift patterns on different microphones.

With tracks aligned to a few seconds and only six false positives in the case study, we are confident that `callsync` is a robust and useful tool for bioacoustics research. Additionally, one false positive was actually a true positive; in this case because of a tiny difference in start and end time between the ground truth and automatic detection, which led the ground truth to be filtered out, but the automatic detection to stay. Overall, the true positive rate of our results was 81%, meaning that only 19% of the manually selected calls for ground truthing were not detected by the `call.detect` function. Where call rate across call types is important, researchers can set the threshold very low and manually remove false positives. Alternatively, a deep neural network can be used to sort signal from noise (Bergler et al. 2022).

A further possible challenge with the `call.detect` function could be that certain call types are never easily distinguishable from background noise. In these situations, `call.detect` is likely to pick up a significant amount of background noise in addition to calls. Function parameters can be adapted and should function on most call-types, as can post-processing thresholding. For example, machine learning approaches (Bergler et al. 2022; Cohen et al. 2022; Stowell et al. 2019) or image recognition tools (Smith-Vidaurre, Araya-Salas, and Wright 2020; Valletta et al. 2017) can be later applied to separate additionally detected noise in particular circumstances where an amplitude based thresholding approach is insufficient. As well, in specific cases (e.g., low amplitude calls), once the align function is performed, the entire `call.detect` function can be swapped out for deep learning call detection algorithms such as ANIMAL-SPOT (Bergler et al. 2022) or other signal processing approaches (e.g., *seewave* package).

The microphones used in this case study were implemented in a captive setting where all calls were within hearing range of each other and each microphone. Thus, all microphones contained a partially shared noisescape. Despite their proximity to one another, it should be noted that each microphone still contained unique vocal attributes, such as wing beats and scratching, and yet the major alignment step still aligned all chunks correctly. This was the case both in recordings with beep markers and without. It is possible that researchers find that the noise differences between microphones is too high for the first alignment step to perform adequately. This would be particularly salient in field settings where individuals within fission-fusion groups find themselves in the proximity of other group members only some of the time (Furmankiewicz et al. 2011; Balsby and Bradbury 2009; Buhrman-Deever, Hobson, and Hobson 2008), or in situations where animals constantly move (i.e., flying) or do independent behaviours that other group members do not (e.g., preening) (Demartsev et al. 2023). Researchers will have to assess their own dataset and test this package to determine whether the first step will perform well on their dataset. If it does not, `callsync` is a modular package and other approaches, such as deep learning (O’shea and West 2016) can be used instead of the first align function, while still using other components of the pipeline. Alternatively, beep markers audible on all microphones could be implemented to assist with alignment. Of course, the audible tones would need to be audible on all microphones.

Assigning a focal bird may be more difficult if individuals tend to be in very close proximity to other group members. If the group members spend time too close to one another (in our case less than 5-10 cm, the distance between the backpack and head of the vocalising individual) (Kloepper and Kinniry 2018; Boughman and Wilkinson 1998), it will become more difficult to distinguish the vocalising individual from other surrounding individuals. In the context of this case study, it was sufficient to only select a vocalising individual when the second loudest call was at least 5% quieter than the focal call. This threshold can be adapted if needed, or the assignment part can be removed entirely if so chosen. More generally, the issue of variation in spatial proximity could be solved by including other on-board or group-level tools, such as accelerometers (Gill et al. 2015; Anisimov et al. 2014), or proximity tags (Wild et al. 2022). This adds extra weight and cost to on-board devices, and so if not possible, the other analytical tools could be considered to distinguish the focal calling birds. The specific tool will depend heavily on use-cases, but include using discriminant analysis (McIlraith and Card 1997) or cepstral coefficients (Lee, Lee, and Huang 2006) to predict the ID of the caller.

Lastly, both fundamental frequency tracing and SPCC work well in certain contexts. For example, automatic fundamental frequency traces work best for tonal calls while SPCC works best when the signal to noise ratio is sufficiently low (Cortopassi and Bradbury 2000). However, if these criteria are not met, several other tools can be used to manually trace fundamental frequency instead, such as Luscinia (Lachlan, Ratmann, and Nowicki 2018) or manual tracing (Araya-Salas and Smith-Vidaurre 2017). While it is important to consider all possible limitations of `callsync` it should also be noted that there are few tools that exist that perform this much needed task. Indeed, the fine scale alignment step of `callsync` allows for call and response dynamics to be measured regardless of how close the calls are to one another. While this paper has thus far only addressed on-board microphones, other systems that implement passive acoustic monitoring systems (Thode et al. 2006) and microphone arrays (Blumstein et al. 2011) should also find benefit within this package depending on the set-up and degree of drift. Possible future research opportunities include trying to incorporate machine learning and noise reduction techniques so that the major alignment can perform in all contexts.

## Conclusion

This open-source package is publicly available on GitHub and CRAN. We welcome all continued suggestions and believe that our package will result in an increase in the scope of bioacoustic research. Our package provides functions that allow alignment, detection, assignment, tracing and analysis of calls in a multi-recorder setting where all microphones are within acoustic distance. The package can be used to generate a fully automated pipeline from raw recordings to the final feature vectors. We show that such a pipeline works well on a captive dataset with four hour long recordings from backpack microphones on six cockatiels which experience non-linear time drift up to several minutes. Each module can also be replaced with alternatives and can be further developed. This package is, to our knowledge, the first R package that performs this task. We hope this package expands the amount of data researchers can process and contributes to understanding the dynamics of animal communication.

## Acknowledgements

We thank Dr. Ariana Strandburg-Peshkin for early feedback. SQS and SAT received funding from the International Max Planck Research School for Organismal Biology and the International Max Planck Research School for Quantitative Behaviour, Ecology and Evolution. SAT received additional funding from a DAAD PhD fellowship. LMA was funded by a Max Planck Group Leader Fellowship from the Max Planck Society, and is currently supported by the Swiss State Secretariat for Education, Research and Innovation (SERI) under contract number MB22.00056.

## Ethics statement

Data from cockatiels were provided by LMA, SAT, and BCK under ethical permission from Regierungspräsidium Freiburg Az. 35-9185.81/G-18/009 within a larger study investigating social learning in cockatiels.

## Data accessibility

All code associated with this paper is publicly available on GitHub (https://github.com/simeonqs/callsync_an_R_package_for_alignment_and_analysis_of_multi-microphone_animal_recordings). All data and code is also publicly available on Edmond (https://edmond.mpdl.mpg.de/privateurl.xhtml?token=05b4aa17-a9bd-44ac-b62b-bdf620aceebb). A DOI for the Edmond repository will be added when the manuscript is accepted and assigned a DOI. The `callsync` package can be installed from CRAN and a developmental version can be found on GitHub (https://github.com/simeonqs/callsync).

## Author contributions

SQS: Conceptualization, Data Curation, Formal Analysis, Methodology, Project Administration, Software, Validation, Visualization, Writing – Original Draft Preparation, Writing – Review & Editing; SAT: Conceptualization, Data Curation, Investigation, Methodology, Project Administration, Validation, Writing – Original Draft Preparation, Writing – Review & Editing; BCK: Investigation, Supervision, Writing – Review & Editing; GAN: Investigation, Methodology, Writing – Review & Editing; LMA: Funding Acquisition, Resources, Supervision, Writing – Review & Editing.

